# Dexmedetomidine attenuates the injury of H9C2 cardiomyocytes under Hypoxia/reoxygenation condition partly through the inhibition of endoplasmic reticulum stress

**DOI:** 10.1101/2020.05.04.076455

**Authors:** Zhipeng Zhu, Xiaoyan Ling, Hongmei Zhou, Caijun Zhang

## Abstract

**Background:** Myocardial ischemia-reperfusion injury (MIRI) has been confirmed to induce endoplasmic reticulum stress(ERS) during downstream cascade reaction when myocardial cell function keep deteriorating to a certain degree. The fact of matter is the clinical inconsistence with experimental outcomes still exist due to the mechanism has not been entirely clarified. Dexmedetomidine (DEX), a new generation anti-inflammatory and organ protector, has been testified can attenuate the IRI of heart. This study aimed to find out if DEX had the capacity to protect the injured cardiomyocytes under in vitro hypoxia/reoxygenation circumstance and if the ERS was totally or partly intervened.

**Methods:** H9C2 cells were subjected to cytotoxicity detection for 24h with DEX normally cultivated in several different concentrations. The proper hypoxia/reoxygenation (H/R) model parameter were concluded by the cell viability and injuries by cell counting kit-8(CCK8) and lactate dehydrogenase (LDH) release, when undergoing hypoxic condition for 3 h and reoxygenated for 3h, 6h,12h, and 24h, respectively. Also, the above index was assessed for H/R cardiomyocytes cultivated by various concentrations of DEX. The apoptosis, expression of) Glucose-regulated protein 78(GRP78), C/EBP homologous protein (CHOP), and caspase-12 were also examined in all groups.

**Results:** 1, 5 and 10 μM DEX in normal culture could significantly promote the proliferation of H9C2 (> 80%); the activity of H9c2 cells decreased to 62.67% (P < 0.05) at 3h of reoxygenation and to 36% at 6h of reoxygenation followed by 3h anoxic treatment; The cell viability of H9c2 cells in H/R groups incubated with 1 μM DEX increased 61.3%, and the LDH concentration in the supernatant was effectively lowered (−13.7, P < 0.05); H/R dramatically decreased the proportion of flow cytometry apoptosis and increased the expression of GRP78, CHOP and caspase-12, while both DEX and 4-phenyl butyric acid (4-PBA) could significantly reverse those above indicators. Additionally, DEX could induce deeper alterations than 4-PBA on the basis of H/R.

**Conclusion:** 1 μM DEX can dramatically attended the cell injuries, apoptosis, the expression of GRP78, CHOP and caspase-12 of H9C2 induced by 3h’ hypoxia and 3h’s reoxygenation. moreover, the functions of DEX went beyond the inhibition of ERS under this situation.

## INTRODUCTION

Myocardial ischemia-reperfusion injury (MIRI), usually happened in clinical settings in an inevitable manner, always confers severe outcomes for patients if no effective strategies to induce the downstream apoptotic cascades advance toward the right direction. Fortunately, numerous animal studies conceiving diverse protective mechanisms have confirmed the efficacy of cardioprotection in conquering the MIRI^[1–6]^. The actual clinical application of part of those strategies, however, turns out to be another scenario, the consequence from clinical practice seems difficult to be consistent with those from experimental research^[7–9]^. In this context, unclear mechanism about the develop of apoptotic pathway and some key molecules throughout MIRI might significantly matter, especially for cell death and even associated signaling pathway during ischemia-reperfusion such as the ERS related apoptosis or signaling pathway.

MIRI is a well-known cause that induce severe damage in endoplasmic reticulum, it was early advocated in Wu’s report in 2016 that the ERS should be taken into consideration wherever MIRI happens^[10]^. In recent years, IRI has been proved to be a multifactorial process that could result in diverse organ damage with the compatible participation of ERS and associated apoptosis. The underlying mechanism consists of excessive oxidative status, ATP depletion and energy imbalance, calcium homeostasis and so on. A growing number of studies have clarified the intervention effect on ERS to the prognosis of MIRI in animal experiment as well as in vitro cell experiment^[11–13]^, and numerous signaling pathways^[14–17]^ have been found involved into ERS such as miR-34a/Sirt1/Nrf2, AMPK/Nrf2, PI3K/AKT, TLR4-MyD88-NF,et al. Furthermore, a noticeable report by Davidson in 2019 hold the opinion that multitarget strategies are necessary to be adopted to reduce MIRI if appropriate, because the single approach has limited capacity to overcome complicated MIRI situation^[18]^.

DEX, as a highly selective *α*^2^ adrenergic receptor agonist, is frequently used in clinical practices, especially provide protective effect on heart as well as other organs throughout operation^[19–22]^. At this point, the most possible function applied could be related to the anti-inflammatory response and inhibition of ischemia reperfusion injury. As for ERS aspect, its effect on ERS and following apoptosis have not been illustrated thoroughly. Judging from those existed studies, it’s noticeable that DEX has shown the protective role on inhibiting IRI to heart of diabetic mice by interfering with ERS or Autophagy^[6,23]^, however, the results of which partly contributed to the diabetes context. Furthermore, researchers put more attention on studying other non-cardiac cell such as endothelial cells under IRI or H/R condition^[24,25]^ as well as examining several crucial ERS chaperones, proteins and apoptosis indicators produced by organs except heart under IRI or H/R condition^[6,26–30]^, finding that DEX can effectively regulate the function of non-cardiac cell and interfere in endoplasmic reticulum stress signal pathway under respective circumstance. There also had few studies explored the function of DEX on H9C2 cardiomyocytes in H/R condition^[31,32]^, the exact regulation function of DEX, however, on the influence on ERS remains unknown, let alone the appropriate experimental condition based on regulation of H/R and if DEX can protect IRI through the inhibition of ERS alone or other multiple function. In this study, we hypothesized that DEX could prevent the cardiac cells from injuring in H/R condition through more than intervention of ERS, trying to verify its capacity to protect the injured cardiomyocytes under in vitro hypoxia/reoxygenation circumstance and optimized the suitable experimental conditions for upcoming research.

## METHODS

### Cell culture

H9C2 cardiomyocytes, rat embryonic myocardial cell line, were obtained from the sample database of central experimental laboratory of the Second Hospital of Jiaxing university. The cells were unfreeze firstly and then cultured in 45g/L glucose DMEM (Corning, USA), with 10% fetal bovine serum (FBS, Invitrogen, USA), 100 g/mL streptomycin and 100 U/mL penicillin (Solarbio, China) added as well. All experimental Cells needed were grown in a 95% air plus 5% CO2 incubator. The medium was replaced according to the growth of cells, which normally was 1–3 days. A 70–80% confluence of cells was ready for experimental use.

### Toxicity test

The original DEX solution was made by DEX powder (SML0956; Sigma-Aldrich; Merck KGaA, Darmstadt, Germany) diluted with DMSO, then the following procedure was carried out and diluted more than 1000 times by DMEM medium, making sure the final concentration of DEX in medium reached 0 nmol / L, 1 nmol / L, 5 nmol / L, 10 nmol / L, 0.1 µmol / L, 1 µmol / L, 5 µmol / L, 10 µmol / L, respectively. All H9C2 cells were incubated in 96-well plates under normal condition overnight with 3 replicates per group, the medium was replaced by another medium full of DEX solution 24h prior to the cell viability test which was conducted by CCK8 kit according to the instructions of manufacturer. Cell viability was represented as relative percentage compared by experimental groups with control group.

### H/R injury model

In order to establish an optimal in vitro H/R model, H9C2 cells, which were made into single cell suspension and incubated on 96 well plate with roughly 5 * 10^3^ / ml cell density in advance, were cultivated in two plates (control plate and experiment plate), with each plate randomly divided into four groups such as H3/R3 group, H3/R6 group, H3/R12 group and H3/R24 group, respectively. The control plate was incubated in 37°C 5%CO_2_ cell incubator and the medium were changed at similar time point with experimental plate. All groups in experimental plate underwent a three-hour anoxia with all cells cultured at 37°C in hypoxia chamber filled with 5%CO_2_ and 95%N_2_ as well as the application of serum-free DMEM when the anoxia procedure began. According to different protocol adopted in the study, those experimental groups received normoxic culture when the reoxygenation procedure started at the end of previous step, the experimental H9C2 cells in respective groups underwent 3hours, 6hours, 12hours, and 24hours’ reoxygenation culture in 37°C 5%CO_2_ cell incubator. At the end of each procedure, the cell viability and damage were detected and tested by CCK8 and LDH kits respectively. The parameters of group that having the best protective properties would be recognized as an optimal condition for H/R group which would be multiple used in following steps. In order to analyses the possible effect of DEX on H9C2 cells under H/R condition, all cells were divided into control group, H/R group and H/R + DEX group. For H/R + DEX group, it included 7 subgroups due to 7 different concentrations of DEX used 1h prior to the start of hypoxia in this study, which was 0 nmol / L, 1 nmol / L, 5 nmol / L, 10 nmol / L, 0.1 µmol / L, 1 µmol / L, 5 µmol / L, 10 µmol / L, respectively. In order to maintain the experimental balance between these groups, the same dose of DMSO that added into other groups was added in control group which was cultured in DMEM supplemented at 37°C with 10% FBS for 6h. The H/R and H/R + DEX group cells were incubated for 3h’s hypoxia in hypoxia chamber filled with 5%CO_2_ and 95%N_2_ followed by 3h’s regeneration under normoxic condition. The LDH concentration of all groups was detected in supernatant and cell viability was detected with CCK-8 kit too.

### Experimental protocols

After accomplishing those above procedures, the optimal dose of DEX for intervention and the best experimental conditions of anoxia/reoxygenation had been built up which would will be adopted by the following experiments. In order to verify that if the present concentration of DEX was an effective condition to attenuate the H/R injuries of H9C2 cells and if the experimental research could be completed on 3h’s hypoxia and 3h’s regeneration model, cultured cells on 96 wells plate and or 6 wells plate were divided into control group, H/R group, H/R+DEX group and H/R+DEX+4-PBA (4-phenylbutyrate, P-21005, Sigma, USA) group. The 4-PBA powder, which is a classic chemical chaperone used to reduce protein misfolding and ameliorate ERS, was dissolved with DMSO solution in sterile and dark environment, and was added into the medium 24h prior to the H/R with the proportion of 1:1000. Apart from the cell viability and injuries tested above, the flow cytometry was adopted to find out the apoptosis of all groups. The last but not least, the expression of GRP78, chop and caspase-12 in terms of protein concentration and mRNA expression were also detected by RT-PCR and Western blot analysis (the specific methods would be described below).

### Cell viability and injury assay

According to the instructions of CCK8 kit (Dojindo, Japan), H9C2 cells (roughly 5*10^3^ cells/well) were seeded into a 96-well plate and followed by associated protocols. When required for viability test, cells in each well were covered with 10 μL Beyotime for at least 1h at 37°C in cultivation incubator. Then the OD value was measured at 450nm with a microplate reader (Molecular Devices, Sunnyvale, CA, USA). The cell viability (%) was calculated thereafter. Meanwhile, the Cell injury was assessed by the amount of LDH release in supernatant with LDH kit (Beyotime Biotechnology, China). According to the LDH assay kit, all cell groups were plated in 96-well plate, with 3 duplications each group, measured with a microplate reader at 450 nm. The LDH concentration and cell injury rate were all calculated.

### Apoptosis assay

The apoptosis rate of H9C2 cells in each group was examined by the flow cytometry with Annexin V-FITC/PI (propidium iodide) kit (Sigma, USA). Briefly, after be digested and centrifuged, cells were washed with PBS twice and suspended with the binding buffer, and then cells incubated with 10 μL Annexin V-FITC as well as 5 μL PI for 20 min at the same time. The apoptosis rate was detected using a flow cytometer (BD, San Jose, CA) and calculated as well.

### Reverse Transcription-Polymerase Chain Reaction (RT-PCR)

The H9C2 cells were prepared for mRNA analysis in advance. The primer sequences of GRP78, caspase12, CHOP, and β-actin were synthesized by Invitrogen company and shown in table 1. The total RNA of H9C2 cells were extracted with Trizol approach, the purity of RNA of each group was determined. With the adaptation of Reverse Transcription Kit (Takara, China), 500ng RNA of each sample was reversely transcribed into cDNA according to the manufacturer’s instructions. Subsequently, the polymerase chain reaction was launched with 2µL cDNA and other necessary ingredients according to the instructions of Premix Ex Taq kit (Takara, China), the ultimate 25µL reaction volume was board on the ABI Prism 7500 System (Bedford, USA). Amplification process initiated at 95 °C for 3min, followed by 40 cycles of amplification, for each cycle including denaturation at 95 °C for 30s, annealing at 55 °C for 20 s, and elongation at 72 °C for 20s. The mRNA expression was concluded through the relative ratio of target gene and β-actin by using the 2 ^-ΔΔCT^ method.

**Table 1.**
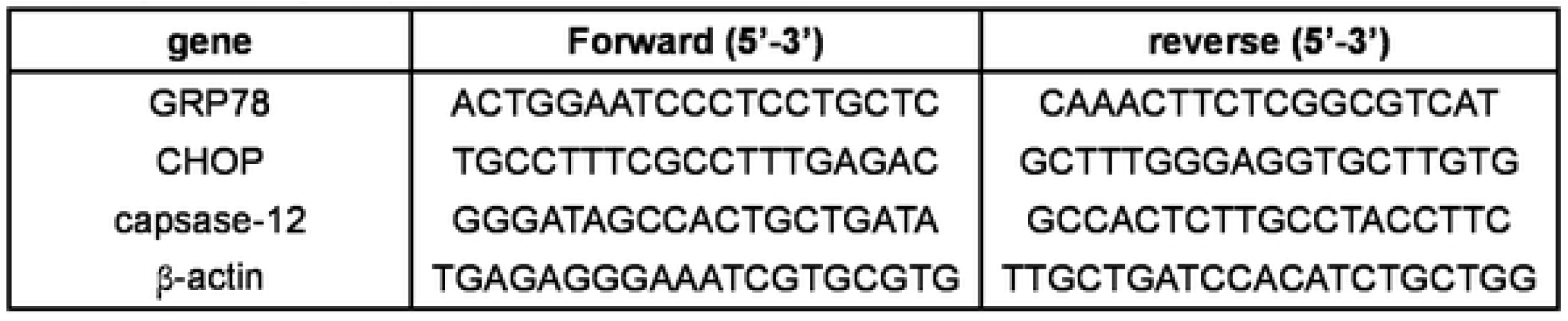
primers oligo sequences.

### Western blots

Following the treatment of previous procedures, each well of the plates culturing H9C2 cells was washed twice to triple with cold PBS solution, and then added RIPA lysates (Beyotime, China) into wells for nearly 30 min on ice. Subsequently, the supernatants were collected after centrifugation of lysates. The BCA method was used to test protein concentrations. The proteins were separated by SDS-PAGE and transferred to PVDF membrane at 4 °C at 200 mA for 2 h. Then, the membranes were blocked in TBST solution for 2 h at room temperature, and incubated at 4 °C overnight with those following primary antibodies: CHOP (Affinity 1: 1000), GRP78 (Affinity 1: 1000), caspase12 (Affinity 1: 1000) and mouse anti-β-actin (Affinity 1: 1000). The second antibody was then combined with horseradish peroxidase (Jackson 1: 2000) for 2 hours at room temperature. Finally, the membrane was cleaned in TBST and the signal was displayed by the enhanced chemiluminescence detection kit (ECL, Beyotime, China). The density of protein bands was quantified by IQuantTL (GE Healthcare, USA). The relative protein expression was calculated with the normalization of β-actin.

### Statistical analysis

All data were expressed with mean ± standard deviation (SD), and One-way analysis of variance (ANOVA) followed by Dunnett’s post hoc test was performed to examine the statistic difference among multiple groups with SPSS 19.0 software (SPSS, Chicago, USA), whereas the Graphpad prism(San Diego, USA) was applied to construct the graphics. A statistically difference was considered in case of P<0.05.

## RESULTS

### DEX promote the proliferation of H9C2

In this study section, a wide range of DEX concentrations which not only involved those be mostly used in clinical setting but those popularly studied in laboratories were chose and put into present study. it was obvious that all those concentrations of DEX selected had no cytotoxicity to H9C2 cells, which represented the higher viability compared to control group (control group marked as 1). However, the 1µM and higher DEX showed extraordinary vitality characteristics with 81% increase for 1µM, 89% for 5µM and 80% for 10µM, compared to control group statistically (Figure 1). According to this result, 1µM, 5µM and 10µM DEX were used in the subsequent experiments.

**Figure 1.**
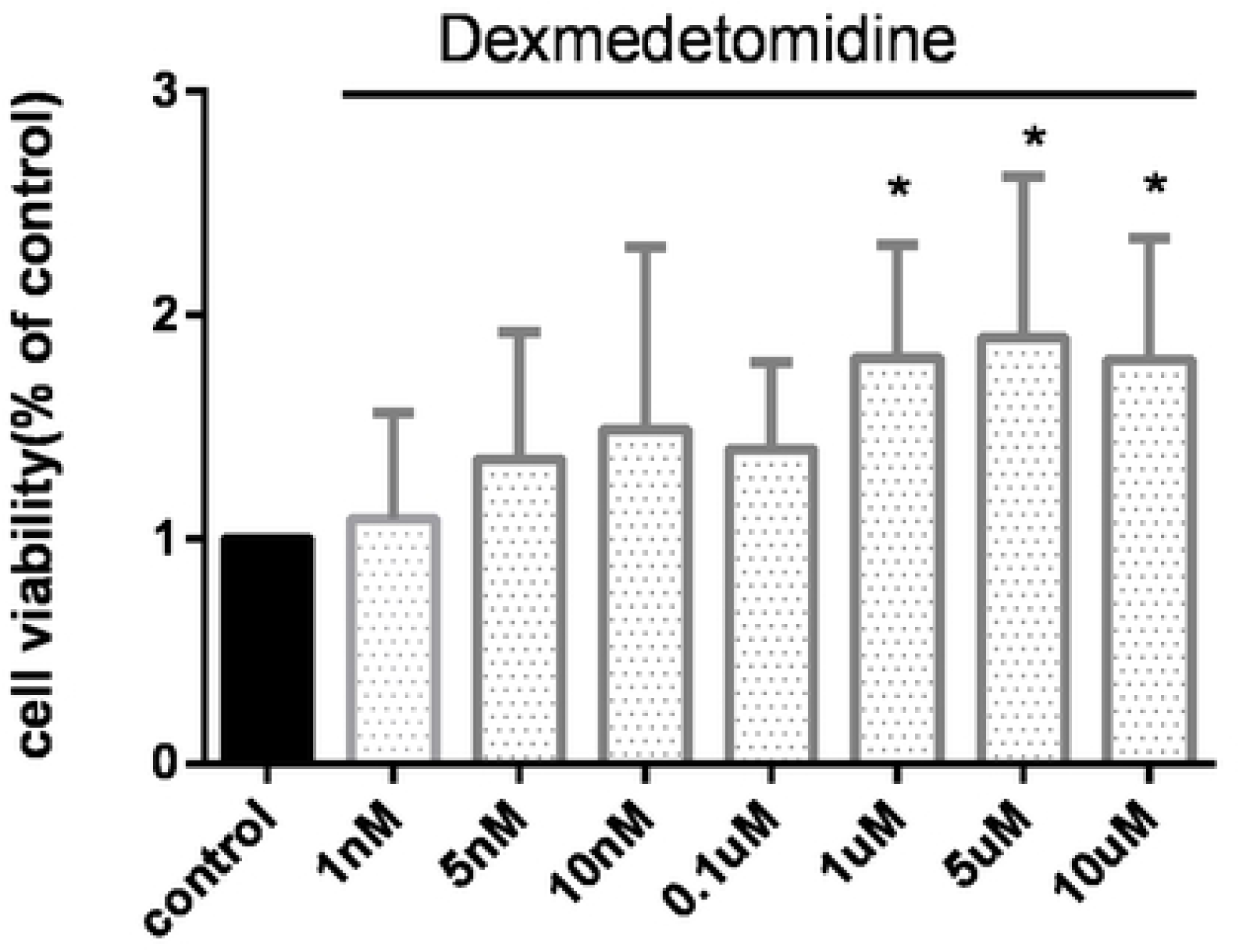
Effects of different concentrations of dexmedetomidine on cardiomyocyte viability when myocardial H9C2 cells were normally cultivated. The results were expressed as mean ± SD. *p < 0.05 vs Control group.

### The 3h Hypoxia / 3h reoxygenation model achieve an optimal experimental level

In order to create the optimal H/R condition to meet the upcoming need, several model parameters were selected such as 3h Hypoxia / 3h reoxygenation, 3h Hypoxia / 6h reoxygenation, 3h Hypoxia / 12h reoxygenation and 3h Hypoxia / 24h reoxygenation (Figure 2.A and B). we investigate the cellular viability and LDH release of H9C2 cells studied on different H/R circumstance. As shown in Figure 2. A, the viability of hypoxia cells tended to decrease at 3h’s reoxygenation, and kept remaining the lower levels until 24h after reoxygenation. Compared to control group, the cell viability in these groups went down by 62.67%, 36%, 53.33% and 58% respectively (P<0.05). In addition, the LDH release assay was used to detect the injuries of H/R H9c2 cells. From Figure 2. B, we found that H/R exposure significantly increased the level of LDH in supernatant, which statistically increased since 3h of reoxygenation (mean =480% of control, P<0.01) until hit the peak at 6h of reoxygenation. All in all, the 3h Hypoxia / 3h reoxygenation model was adoptable for following experiments.

**Figure 2.**
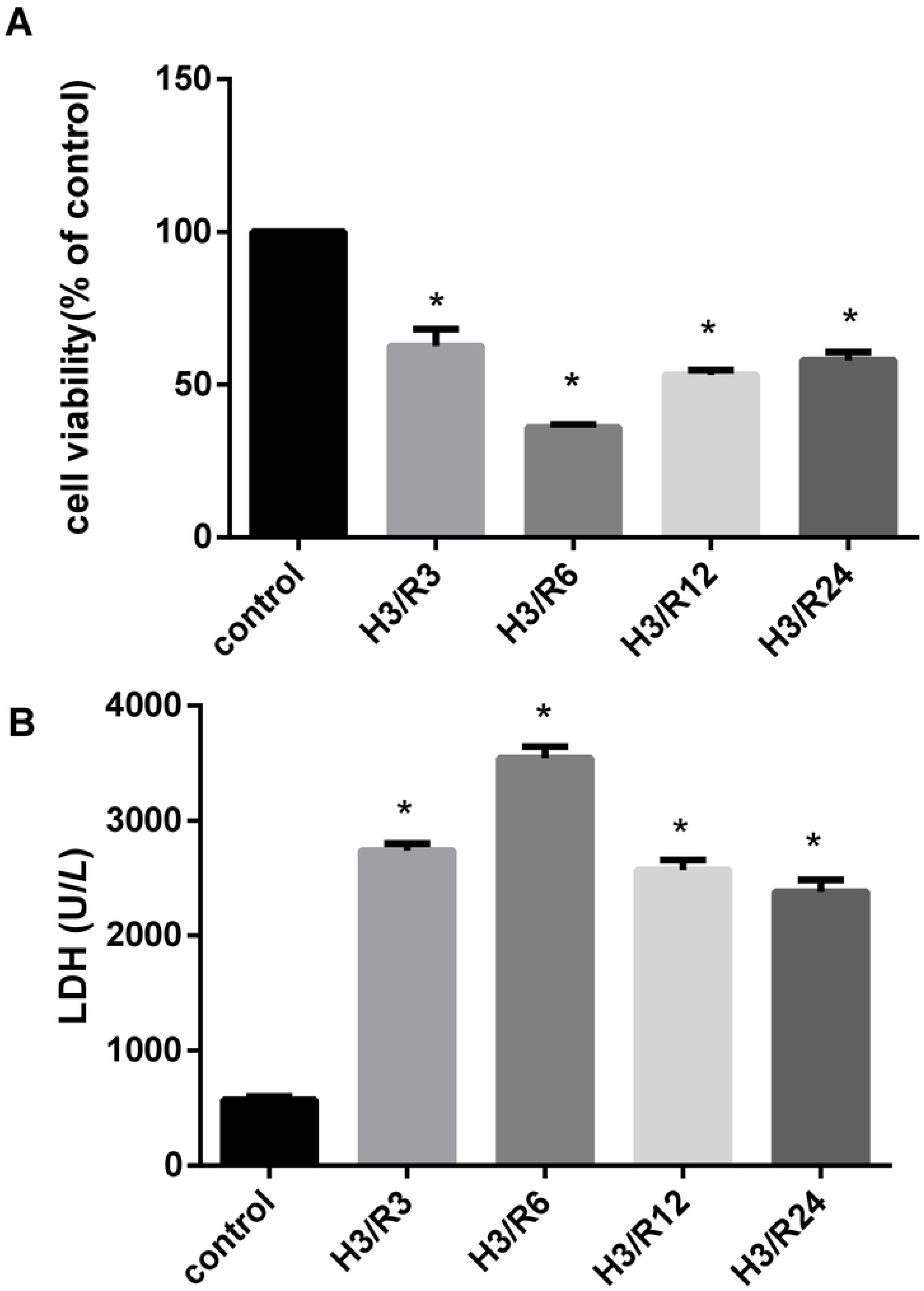
levels of myocardial cell injuries due to H/R under different strategies are illustrated in (A)proportions of cell viability and (B) LDH. The results were expressed as mean ± SD. *p < 0.05 vs Control group. LDH, lactate dehydrogenase; H, hypoxia; R, reoxygenation.

### 1µM DEX could effectively protect H9C2 cells from H/R injury

Based on those above H/R experimental results, we investigated the influence of DEX on H9C2 injuries on the ground of 3h Hypoxia / 3h reoxygenation. As for concentration of DEX applied in this section, those that showed the most effectiveness in promoting the proliferation of H9C2 such as 1µM, 5µM and 10µM were adopted to test the viability and injuries of H9C2 cells during H/R. As shown in Figure 3, the three concentrations of DEX could all level up the viability of H/R H9C2 cells, however, it showed gradually less effect from 1µM to 10µM, and the most functional group was 1µM group compared to control group. Similarly, the 1µM DEX pretreatment to H/R H9C2 cells obviously inhibited the LDH release by 113.7% compared with control group (P<0.05).

**Figure 3.**
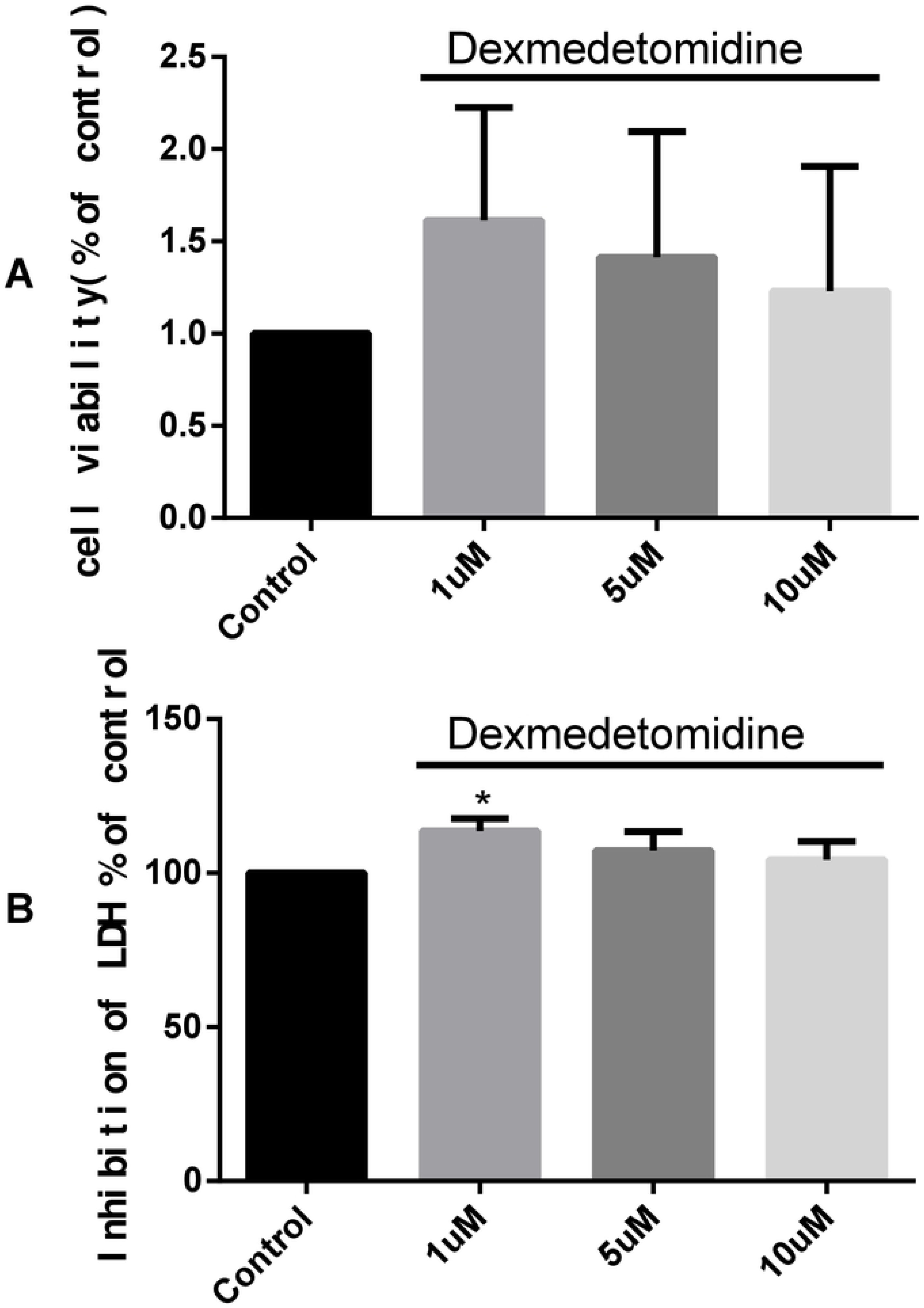
Effects of DEX of different incubation concentration on levels of myocardial cell injuries due to H/R are illustrated in (A) proportions of cell viability and (B)the percentage of the inhibition of LDH. The results were expressed as mean ± SD. *p < 0.05 vs Control group. LDH, lactate dehydrogenase; H, hypoxia; R, reoxygenation; DEX, Dexmedetomidine.

### DEX reduced the injuries of H9C2 cells as well as apoptosis

In order to elaborately investigate the function of DEX on H9C2 cells during H/R, several interfering strategies were applied in this branch experiment (Figure 4 and 5). When taking into account the cell viability and LDH release, the control group, the DEX pretreatment group under normal condition and the 4-PBA pretreatment group under normal condition saw the similar comparable changes (P>0.05). However, it was noticeable that the H/R group showed dramatic decrease in terms of cell viability and high amount of LDH release compared with the control group (P<0.05) which identically corresponded to the previous results. Compared with H/R group, 1µM DEX could alleviate the cellular injury furtherly as much as the 4-PBA did. In addition, the combination of DEX and 4-PBA which described in DEX+4-PBA+H/R group showed an advancing protective effect on cell viability than the sole group respectively (Figure 4. A and B). As for the cell apoptosis assay, there was an average 10% decline in H/R group compared with control group (Figure 5). Similarly, DEX+ H/R or 4-PBA+ H/R pretreatment group had more severe cell apoptosis which accounted for higher percentage by 15% or 19% than H/R group (P<0.05). The combination of DEX and 4-PBA showed more beneficial function than single intervene as well (Figure 5).

**Figure 4.**
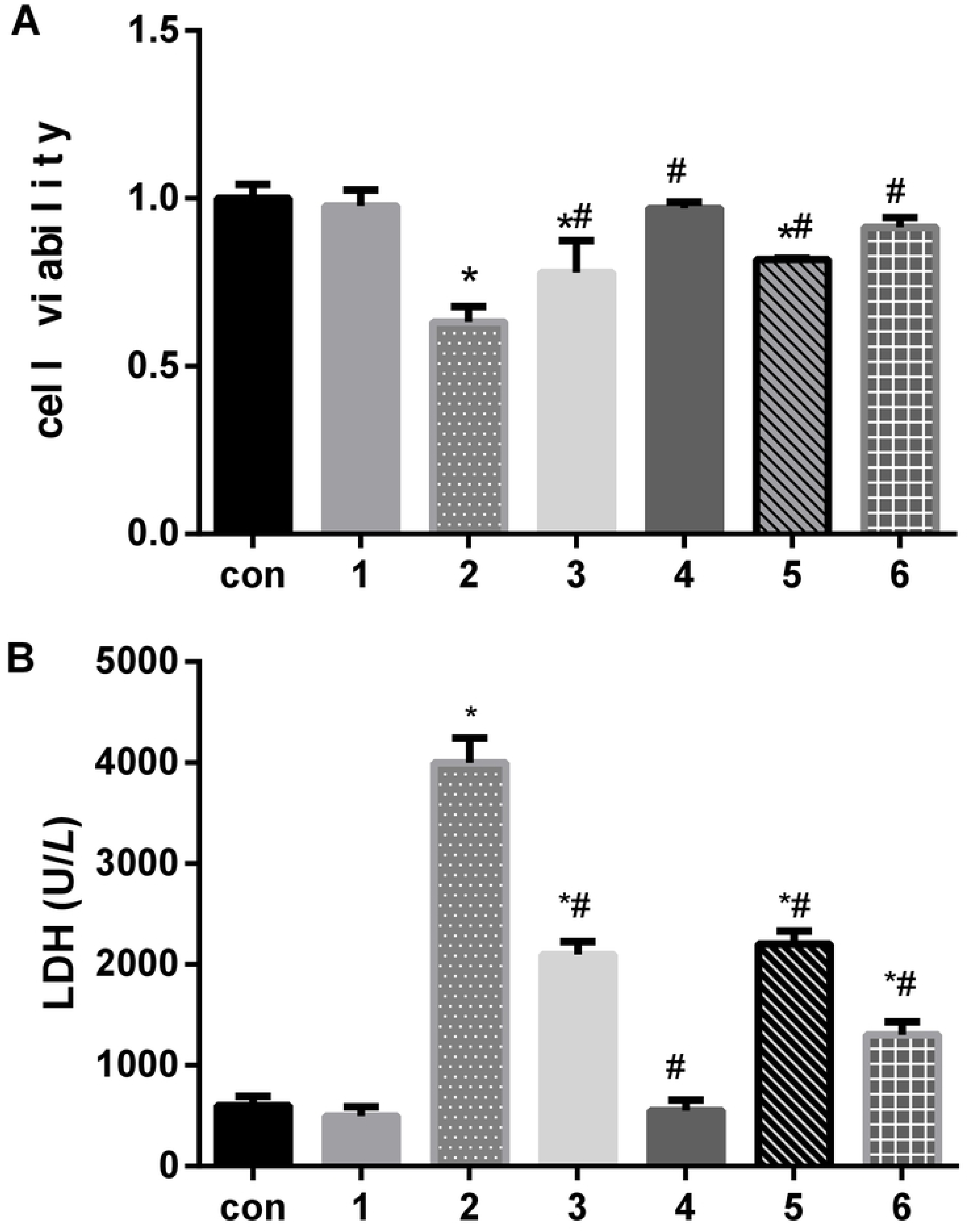
Effects of dexmedetomidine on levels of myocardial cell injuries due to H/R under different strategies are illustrated in (A)proportions of cell viability and (B) LDH. The results were expressed as mean ± SD. *p < 0.05 vs Control group; #P<0.05 VS H/R group. con,control group;1,normoxia incubation with dexmedetomidine,2, hypoxia/reoxygenation incubation;3,hypoxia/reoxygenation incubation with dexmedetomidine;4. normoxia incubation with 4-phenyl butyric acid;5,hypoxia/reoxygenation incubation with 4-phenyl butyric acid;6.hypoxia/reoxy genation incubation with dexmedetomidine4-phenyl butyric acid. LDH, lactate dehydrogenase; H, hypoxia; R, reoxygenation.

**Figure 5.**
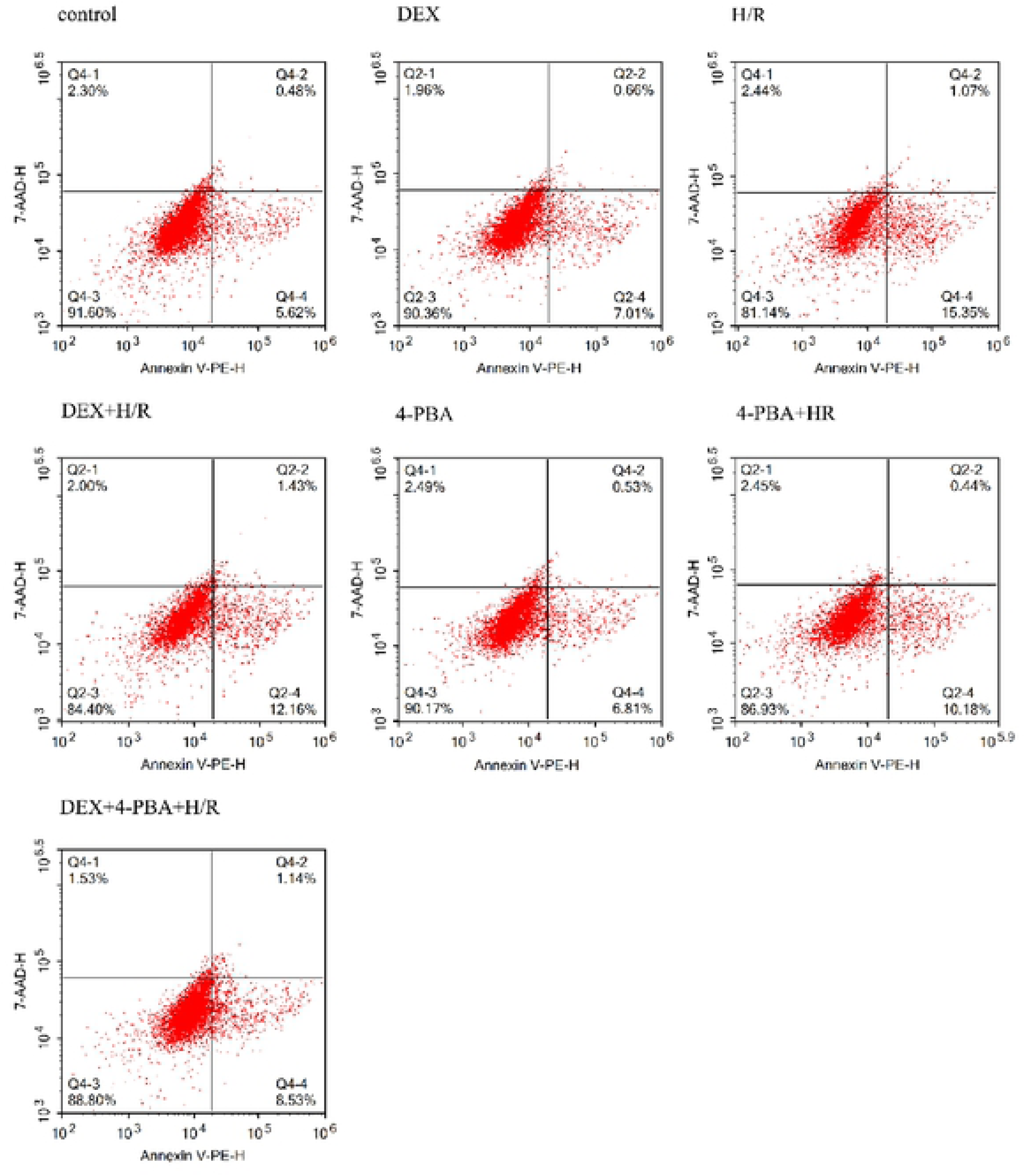
The apoptosis percentages of H9C2 cardiomyocyte were examined by flow cytometry under different strategies. DEX, dexmedetomidine; 4-PBA, 4-phenyl butyric acid; H/R, hypoxia/reoxygenation.

### DEX reduced the expressions of GRP78, CHOP and Caspase-12 in H9C2 cells after H/R

With the goal of figuring out if DEX was involved with those above function, three main molecules closely attached to ERS and related signing pathway were examined (Figure 6). As illustrated in Figure 6, consistent results were reached between mRNA expressions and protein expressions in similar experimental design as previous. Compared to the control group, H/R group saw obvious upregulation of indicators in terms of ERS and apoptosis. DEX or 4-PBA pretreatment based on H/R condition significantly alleviated the expressions of GRP78, CHOP and Caspase-12 respectively (P<0.05). Moreover, a more decreasing decline was also found in the combination group such as DEX+ 4-PBA+ H/R, compared to DEX+ H/R or 4-PBA+ H/R group which was lonely interfered.

**Figure 6.**
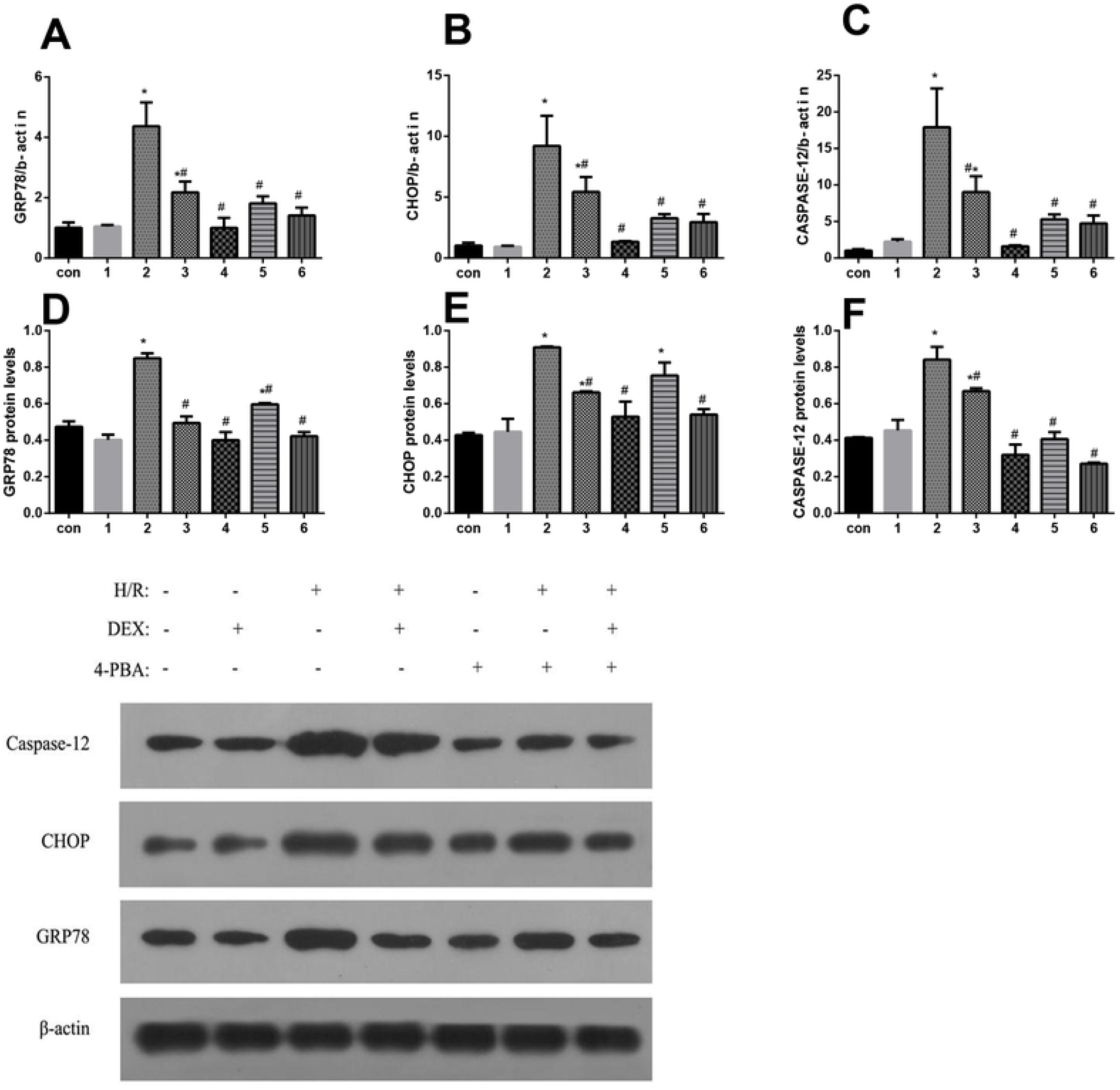
Effects of dexmedetomidine on expression of GRP78, CHOP and caspase-12 when myocardial cells are under different circumstance. A-C, the mRNA expression of GRP78, CHOP and caspase-12; D-F, the protein expression of GRP78, CHOP and caspase-12. The results were expressed as mean±SD. *P<0.05 vs control group; #P<0.05 vs H/R group. Con, control group; 1, normaoxia incubation with dexmedetomidine, 2, hypoxia/reoxygenation incubation; 3, hypoxia/reoxygenation incubation with dexmedetomidine; 4, normoxia incubation with 4-phenylbutyric acid; 5, hypoxia/reoxygenation incubation with 4-phenylbutyric acid; 6, hypoxia/reoxygenation incubation with dexmedetomidine and 4-phenylbutyric acid.

## DISCUSSION

It’s known from previous researches that DEX has myocardial protection under IRI or H/R condition. Presently, with an attempt to elaborately figure out the precise connection between ERS and DEX application under those complicated situation, we built up the classic H/R model in present experiment and exerted common-use tests such as CCK8, LDH, flow cytometry, RT-PCR and western-blot, trying to find out DEX’s intervention with H/R injuries and if the ERS was involved in this process. Also, the optimal experimental condition associated with those procedures were detected. Eventually, this study confirmed that DEX attenuated the myocardial damage brought by H/R through the downregulation of several key molecules relating to ERS and apoptosis, especially the extended cardioprotection beyond the inhibition of ERS.

As a popular phenomenon in clinical setting, the MIRI has correlation with a variety of process. Until now, no effective therapy applied clinically could be used to reduce MIRI, even some of which had been confirmed to be successful in experimental projects such as the use of cyclosporine A^[33]^. Of course, some drawbacks of animal experiments except large animals were underlined for this dilemma due to numerous diversities between animal cardiac physiology and humans. The further clarification of protective mechanism for diverse approaches is another reason that really matters^[34]^, which could be fulfilled by perfectly constructed in vitro models with isolated cardiomyocytes to independently control those external factors^[35]^. So, it’s crucial to reach conclusion from validated in vitro models and makes it clear for important mechanism. Among countless studies in recent years, the necessary hypoxia and reoxygenation time varied from one study to another, which directly lead to the controversy of strategic practice. In addition, the composition of culture media at point of reoxygenation is also a major factor that should take into consideration of nutrients, extracellular pH, calcium concentration, et al^[35]^. It’s stated that 30 min is enough for animal experiment in terms of IRI condition^[36]^, however, there isn’t a consistent time parameters for cardiomyocytes because of the different levels of maturity and oxygen dependency for adopted cells or the cell source used. Xie’s study^[37]^ found that it’s better to adjust the ischemic time in the range of two to five hours. For cardiomyocytes aspect, several cell types, including neonatal cardiomyocytes, adult cardiomyocytes, immortalized cell lines and human pluripotent stem cell, could be chose for experiment. Compared from them, H9c2, as one of immortalized cell, is the most suitable cardiomyocytes for researcher to conduct IRI and toxicology experiments presently if no cellular contract is considered^[38,39]^. Also, The European Society of Cardiology Working Group Cellular Biology of the Heart has clarified that the optimal duration for combined ischemic and reperfusion should be the time that could result in 50% cell death^[40]^, but it is not too long to affect possible intervention effect. Based on all above considerations, we used H9C2 to conduct 3 hours’ hypoxia experiment which was already used by other researchers^[31,41]^ and then several reoxygenation protocols were examined, the roughly 50% cell death was gained at 3 hours’ reoxygenation, although it hit the bottom at 6 hours with 64% decrease.

In recent years, the increasing number of studies have been conducted to testify the protective cardiofunction of DEX on the context of pretreatment or postconditioning^[1,6,24,32]^. Several animal studies have identified that DEX have myocardial protective effect through reducing the ERS after myocardial IRI or regulate the process of myocardial apoptosis which consist of intrinsic and extrinsic apoptotic pathways^[1,6,24,42]^. With an attempt to furtherly understand the mechanism under IRI situation, few cellular level experiments^[17,31,32,43]^ were carried out with different cardiomyocytes or hypoxia / reoxygenation protocols, concluded that DEX could intervene with calcium overload, small non-coding RNAs, inflammation, et al. In our present research, we adopted a wide range of concentration of DEX from 1nM to 10µM for pretreatment to create an optimal experimental condition, directly test those specialized indicator of ERS pathway such as GRP78, CHOP, caspase-12. As far as we know, it’s the first time to examine the protective function of DEX with such a broad range. According to KE’s study^[43]^, the concentration of DEX above 30µM could damage the viability of cardiomyocytes with a certain cytotoxicity, so we selected 10µM as top of the concentration. Our result indicated that DEX showed no cytotoxicity to H9C2 in whatever concentration under 10µM, on the contrary, an obvious cell proliferation was found at higher concentrations such as 1, 5 and 10µM. Consequently, 1µM DEX attenuated the cellular injury of H9C2 induced by hypoxia/reoxygenation, which was similar to other studies conducted by Mei and Gao who used the same dose of DEX^[32]^, but non-ERS pathway were involved.

It has been well accepted that MIRI could lead to severe ERS which was labeled by upregulated GRP78^[44,45]^. If the ERS went deeper, apoptosis cascades would be considered as an underlying mechanism of MIRI, and some specific transcription molecules would be upregulated in an ERS-dependent manner such as CHOP and caspase-12. For them, most researchers confirmed that they can represent the status of ERS and even the developing direction of cell survival as the downstream marker of ERS signaling pathway^[46–48]^. Moreover, ERS could be alleviated in MIRI through the suppression of caspase-12 and CHOP. In this preclinical study, these 3 above molecules were highly expressed under H/R situation, otherwise dramatically alleviated by DEX and 4-PBA. This result thoroughly indicated that DEX showed the potential to involve into ERS-associated apoptosis and had the capacity to inhibit ERS by a similar function of 4-PBA, as reflected by the decreased expression of CHOP, GRP78 and caspase-12 at both mRNA and protein level. Additionally, it should be noticed that DEX exerted more protection against RES-related apoptosis than the inhibition of ERS in DEX+H/R+4-PBA group when compared to H/R+4-PBA group, which indicated a lower expression of CHOP, GRP78 and caspase-12. It is supposed that DEX might intervene with mitochondria-dependent or death receptor-dependent apoptosis except for ERS-associated apoptotic signaling pathway due to unknown reason. In this regard, it’s compatible with Davidson’s opinion^[18]^ more or less that DEX might have multifunction to cope with IRI. More studies should be done in the future focusing on what other function of DEX will be involved this procedure.

There was no denying that several limitations should be taken into consideration. Primarily, this study was just a pretreatment experiment without connection to any signaling pathway, with an obvious attempt to complete some fundamental research for future experiments. Secondly, we admitted that the purpose of any in vitro H/R model is to mimic the real IRI happening in clinical setting, the condition-dependent models might vary from one to another. Furthermore, human cells or stem cells which can more accurately reflect the clinical scene was not investigated in present study. The last but not least, the present study is just a preconditioning investigation about DEX, the effects of postconditioning which is getting a popularization in clinical trial was not in present researching territory. In addition, the further investigation into the precise mechanisms that found in this study should be carried out.

## CONCLUSIONS

As a promising agent in the clinical setting, DEX has been confirmed presently in cellular level that can successfully protect H9C2 cells with a dose of 1 µM under the condition of 3h’ hypoxia and 3h’s reoxygenation. Furthermore, the results also indicated that the functions of DEX could be realized through the intervention of ERS and following apoptosis which were illustrated as the expression of GRP78, CHOP and caspase-12. Besides, DEX showed more advanced protection than the inhibition of ERS by 4-PBA under H/R situation. At this point, more following studies are needed to find out the additional function of DEX in this circumstance and which signaling pathway is involved.

## ACKNOWLEDGMENTS

Thanks to the contribution of all colleagues in the anesthesiology department, the present study has eventually accomplished with and the funding of the Scientific and Technological Projects of Zhejiang Province (NO 2017C33185) and the Project of Health Commission of Zhejiang Province (NO 2019ZD053).

## REFERENCES

1. Bunte S, Behmenburg F, Majewski N, et al. Characteristics of Dexmedetomidine Postconditioning in the Field of Myocardial Ischemia-Reperfusion Injury. Anesth Analg. Jan 2020;130(1):90–98.

2. Li L, Li X, Zhang Z, et al. Protective Mechanism and Clinical Application of Hydrogen in Myocardial Ischemia-reperfusion Injury. Pakistan journal of biological sciences: PJBS. 2020;23(2):103–112.

3. Xi J, Li QQ, Li BQ, et al. miR-155 inhibition represents a potential valuable regulator in mitigating myocardial hypoxia/reoxygenation injury through targeting BAG5 and MAPK/JNK signaling. Molecular medicine reports. 2020; 21(3):1011–1020.

4. Li J, Zhou W, Chen W, et al. Mechanism of the hypoxia inducible factor 1/hypoxic response element pathway in rat myocardial ischemia/diazoxide post-conditioning. Molecular medicine reports. 2020; 21(3):1527–1536.

5. Kitazume-Taneike R, Taneike M, Omiya S, et al. Ablation of Toll-like receptor 9 attenuates myocardial ischemia/reperfusion injury in mice. Biochemical and biophysical research communications. 2019(3):442–447.

6. Li J, Zhao Y, Zhou N, et al. Dexmedetomidine Attenuates Myocardial Ischemia-Reperfusion Injury in Diabetes Mellitus by Inhibiting Endoplasmic Reticulum Stress. Journal of diabetes research. 2019;30:7869318.

7. Heusch G, Gersh BJ. The pathophysiology of acute myocardial infarction and strategies of protection beyond reperfusion: a continual challenge. European heart journal. 2017(11):774–784.

8. Heusch G. Cardioprotection research must leave its comfort zone. European heart journal. 2018(36):3393–3395.

9. Heusch G. Critical Issues for the Translation of Cardioprotection. Circulation research. 2017(9):1477–1486.

10. Wu H, Ye M, Yang J, et al. Endoplasmic reticulum stress-induced apoptosis: A possible role in myocardial ischemia-reperfusion injury. Int J Cardiol. Apr 1 2016;208:65–66.

11. Li W, Li W, Leng Y, et al. Ferroptosis Is Involved in Diabetes Myocardial Ischemia/Reperfusion Injury Through Endoplasmic Reticulum Stress. DNA Cell Biol. Feb 2020;39(2):210–225.

12. Gao J, Guo Y, Liu Y, et al. Protective effect of FBXL10 in myocardial ischemia reperfusion injury via inhibiting endoplasmic reticulum stress. Respir Med. Jan 2020;161:105852.

13. Guo C, Zhang J, Zhang P, et al. Ginkgolide B ameliorates myocardial ischemia reperfusion injury in rats via inhibiting endoplasmic reticulum stress. Drug Des Devel Ther. 2019;26(13):767–774.

14. Wang X, Yuan B, Cheng B, et al. Crocin Alleviates Myocardial Ischemia/Reperfusion-Induced Endoplasmic Reticulum Stress via Regulation of miR-34a/Sirt1/Nrf2 Pathway. Shock. Jan 2019;51(1):123–130.

15. Hou X, Fu M, Cheng B, et al. Galanthamine improves myocardial ischemia-reperfusion-induced cardiac dysfunction, endoplasmic reticulum stress-related apoptosis, and myocardial fibrosis by suppressing AMPK/Nrf2 pathway in rats. Ann Transl Med. Nov 2019;7(22):634.

16. Zhang BF, Jiang H, Chen J, et al. Nobiletin ameliorates myocardial ischemia and reperfusion injury by attenuating endoplasmic reticulum stress-associated apoptosis through regulation of the PI3K/AKT signal pathway. Int Immunopharmacol. Aug 2019;73:98–107.

17. Gao JM, Meng XW, Zhang J, et al. Dexmedetomidine Protects Cardiomyocytes against Hypoxia/Reoxygenation Injury by Suppressing TLR4-MyD88-NF-kappaB Signaling. Biomed Res Int. 20172017:1674613.

18. Davidson SM, Ferdinandy P, Andreadou I, et al. Multitarget Strategies to Reduce Myocardial Ischemia/Reperfusion Injury: JACC Review Topic of the Week. J Am Coll Cardiol. Jan 8 2019;73(1):89–99.

19. Kong Q, Wu X, Qiu Z, et al. Protective Effect of Dexmedetomidine on Acute Lung Injury via the Upregulation of Tumour Necrosis Factor-α-Induced Protein-8-like 2 in Septic Mice. Inflammation. 2020;11: doi: 10.1007/s10753-019-01169-w.

20. Zhang Y, Liu M, Yang Y, et al. Dexmedetomidine exerts a protective effect on ischemia-reperfusion injury after hepatectomy: A prospective, randomized, controlled study. Journal of clinical anesthesia. 2019;5(61): 109631.

21. Xiong J, Quan J, Qin C, et al. Dexmedetomidine Exerts Brain-Protective Effects Under Cardiopulmonary Bypass Through Inhibiting the Janus Kinase 2/Signal Transducers and Activators of Transcription 3 Pathway. Journal of interferon & cytokine research: the official journal of the International Society for Interferon and Cytokine Research. 2019; 40(2):116–124.

22. Gong J, Zhang R, Shen L, et al. The brain protective effect of dexmedetomidine during surgery for paediatric patients with congenital heart disease. The Journal of international medical research. 2019(4):1677–1684.

23. Oh JE, Jun JH, Hwang HJ, et al. Dexmedetomidine restores autophagy and cardiac dysfunction in rats with streptozotocin-induced diabetes mellitus. Acta diabetologica. 2019(1):105–114.

24. He L, Hao S, Wang Y, et al. Dexmedetomidine preconditioning attenuates ischemia/reperfusion injury in isolated rat hearts with endothelial dysfunction. Biomedicine & pharmacotherapy = Biomédecine & pharmacothérapie. 2019:108837.

25. Riquelme JA, Westermeier F, Hall AR, et al. Dexmedetomidine protects the heart against ischemia-reperfusion injury by an endothelial eNOS/NO dependent mechanism. Pharmacological research. 2016;103:318–327.

26. Liu Y, Wang S, Wang Z, et al. Dexmedetomidine Alleviated Endoplasmic Reticulum Stress via Inducing ER-phagy in the Spinal Cord of Neuropathic Pain Model. Frontiers in neuroscience. 2020;28(14):90.

27. Chai Y, Zhu K, Li C, et al. Dexmedetomidine alleviates cisplatin-induced acute kidney injury by attenuating endoplasmic reticulum stress-induced apoptosis via the α2AR/PI3K/AKT pathway. Molecular medicine reports. 2020;21(3):1597–1605.

28. Sun D, Wang J, Liu X, et al. Dexmedetomidine attenuates endoplasmic reticulum stress-induced apoptosis and improves neuronal function after traumatic brain injury in mice. Brain research. 2020; 1(1732):146682.

29. Zhao L, Zhai M, Yang X, et al. Dexmedetomidine attenuates neuronal injury after spinal cord ischaemia-reperfusion injury by targeting the CNPY2-endoplasmic reticulum stress signalling. Journal of cellular and molecular medicine. 2019;23(12):8173–8183.

30. Liu C, Fu Q, Mu R, et al. Dexmedetomidine alleviates cerebral ischemia-reperfusion injury by inhibiting endoplasmic reticulum stress dependent apoptosis through the PERK-CHOP-Caspase-11 pathway. Brain research. 2018:246–254.

31. Wang Z, Yang Y, Xiong W, et al. Dexmedetomidine protects H9C2 against hypoxia/reoxygenation injury through miR-208b-3p/Med13/Wnt signaling pathway axis. Biomedicine & pharmacotherapy = Biomédecine & pharmacothérapie. 2020:110001.

32. Yuan M, Meng XW, Ma J, et al. Dexmedetomidine protects H9c2 cardiomyocytes against oxygen-glucose deprivation/reoxygenation-induced intracellular calcium overload and apoptosis through regulating FKBP12.6/RyR2 signaling. Drug design, development and therapy. 2019;2(13):3137–3149.

33. Cung TT, Morel O, Cayla G, et al. Cyclosporine before PCI in Patients with Acute Myocardial Infarction. The New England journal of medicine. 2015(11):1021–1031.

34. Rossello X, Yellon DM. Cardioprotection: The Disconnect Between Bench and Bedside. Circulation. 2016(8):574–575.

35. Chen T, Vunjak-Novakovic G. In vitro Models of Ischemia-Reperfusion Injury. Regen Eng Transl Med. Sep 2018;4(3):142–153.

36. He X, Li S, Liu B, et al. Major contribution of the 3/6/7 class of TRPC channels to myocardial ischemia/reperfusion and cellular hypoxia/reoxygenation injuries. Proceedings of the National Academy of Sciences of the United States of America. 2017(23):E4582–E4591.

37. Xie M, Kong Y, Tan W, et al. Histone deacetylase inhibition blunts ischemia/reperfusion injury by inducing cardiomyocyte autophagy. Circulation. 2014(10):1139–1151.

38. Oh JG, Kho C, Hajjar RJ, et al. Experimental models of cardiac physiology and pathology. Heart Fail Rev. Jul 2019;24(4):601–615.

39. Kuznetsov AV, Javadov S, Sickinger S, et al. H9c2 and HL-1 cells demonstrate distinct features of energy metabolism, mitochondrial function and sensitivity to hypoxia-reoxygenation. Biochimica et biophysica acta. 2015(2):276–284.

40. Lecour S, Bøtker HE, Condorelli G, et al. ESC working group cellular biology of the heart: position paper: improving the preclinical assessment of novel cardioprotective therapies. Cardiovascular research. 2014(3):399–411.

41. He S, Wang X, Zhong Y, et al. Hesperetin post-treatment prevents rat cardiomyocytes from hypoxia/reoxygenation injury in vitro via activating PI3K/Akt signaling pathway. Biomedicine & pharmacotherapy = Biomédecine & pharmacothérapie. 2017:1106–1112.

42. Yang YF, Peng K, Liu H, et al. Dexmedetomidine preconditioning for myocardial protection in ischaemia-reperfusion injury in rats by downregulation of the high mobility group box 1-toll-like receptor 4-nuclear factor κB signalling pathway. Clinical and experimental pharmacology & physiology. 2017(3):353–361.

43. Peng K, Qiu Y, Li J, et al. Dexmedetomidine attenuates hypoxia/reoxygenation injury in primary neonatal rat cardiomyocytes. Exp Ther Med. Jul 2017;14(1):689–695.

44. Wang R, Yang M, Wang M, et al. Total Saponins of Aralia Elata (Miq) Seem Alleviate Calcium Homeostasis Imbalance and Endoplasmic Reticulum Stress-Related Apoptosis Induced by Myocardial Ischemia/Reperfusion Injury. Cellular physiology and biochemistry: international journal of experimental cellular physiology, biochemistry, and pharmacology. 2018(1):28–40.

45. Bi X, Zhang G, Wang X, et al. Endoplasmic Reticulum Chaperone GRP78 Protects Heart From Ischemia/Reperfusion Injury Through Akt Activation. Circulation research. 2018(11):1545–1554.

46. Li H, Chen H, Li R, et al. Cucurbitacin I induces cancer cell death through the endoplasmic reticulum stress pathway. Journal of cellular biochemistry. 2018;11,doi: 10.1002/jcb.27570.

47. Huang ZH, Zhang SX, Wang C, et al. Downregulated long non-coding RNA FOXD3-AS1 promotes endoplasmic reticulum stress-induced apoptosis by inhibiting RCN1 via let-7e-5p in nasopharyngeal carcinoma. American journal of physiology. Cell physiology. 2020;19, doi: 10.1152/ajpcell.00352.2019.

48. Chen J, Chen J, Cheng Y, et al. Mesenchymal stem cell-derived exosomes protect beta cells against hypoxia-induced apoptosis via miR-21 by alleviating ER stress and inhibiting p38 MAPK phosphorylation. Stem cell research & therapy. 2020;11(1):97.

